# Multisensory perception, verbal, visuo-spatial, and motor working memory modulation after a single open- or closed-skill exercise session in children

**DOI:** 10.1101/2020.01.29.924563

**Authors:** Jessica O’Brien, Giovanni Ottoboni, Alessia Tessari, Annalisa Setti

## Abstract

Physical activity presents clear benefits for children’s cognition; this study examined the effect of a single exercise session of open- or closed-skill exercise, as opposed to a no-exercise activity on multisensory perception, i.e. the ability to appropriately merge inputs from different sensory modalities, and on working memory (verbal, visuo-spatial, and motor working memory) in 51 preadolescent children (aged 6-8). Using a semi-randomised pre-post design, participants completed a range of cognitive tasks immediately before and after an exercise session or a classroom sedentary activity. Participants were randomised, within each school, to one of the three groups (open-skill, n=16; closed-skill, n=16; classroom activity, n=19). Exercise, but not usual classroom activity, improved children’s multisensory perception, with no difference between exercise types. Results also revealed that a single open-skill session produced verbal working memory (digit span) benefits; a closed-skill exercise session benefitted motor working memory. While the relatively small number of participants should be acknowledged as limitation, these findings contribute to emerging evidence for selective cognitive benefits of exercise, and show, for the first time in children, that multisensory processing sensitivity is improved by exercise.

## Introduction

The evidence for the positive effects of physical activity on cognitive functioning (Hillman et al. 2008) coincides with a time of global concern regarding the increasingly sedentary lifestyle of children (Karnik and Kanekar 2012).

Acute exercise (i.e. a single exercise bout) represents an intervention strategy to promote children’s cognitive development (Diamond and Lee 2011), as single exercise sessions can easily be implemented into children’s daily lives, especially within school (Masini et al. 2019). Yet there is a dearth of research on acute exercise on children (Ellemberg and St-Louis-Deschênes 2010) and questions remain open on how to inform practical guidelines on the optimal exercises for cognitive enhancement based on the current knowledge of brain benefits of exercise (Hötting and Röder 2013).

Different exercise types have differential effects on the brain, with emerging experimental evidence indicating cognitively-stimulating exercise (particularly open-skill exercise) may represent the optimal exercise type to improve cognition (Pesce 2009). Open-skill exercise is defined by an unpredictable exercise environment (e.g. team sports) requiring the constant updating of sensory information, movement and attention, for the purpose of appropriate action. Open-skill exercise is therefore characterised by high cognitive demands. This contrasts with closed-skill exercise, characterised by a predictable environment (e.g. running, swimming) and lower cognitive demands (Pesce 2009). Findings in adults document differential neural activation in open- and closed-skill exercisers, with open-skill exercisers displaying better neural efficiency during cognitive tasks (Huang et al. 2014). Importantly, a recent meta-analysis (Gu et al. 2019) highlighted that only few studies directly test exercise in children comparing open- and closed-skill exercise types. In the meta-analysis, which included participants across the lifespan, it was concluded that there is evidence for larger benefits of open-skill exercise over closed-skill exercise in children and older adults, particularly on executive functions. However, this evidence is weakened by the very small amount of studies addressing the topic.

In the present study, we aim to extend to school aged children our findings of a benefit of one bout of open skill exercise on multisensory processing, i.e. the ability to appropriately integrate information from different sensory modalities, and working memory (WM) (O’Brien et al. 2017). These cognitive abilities are foundational to learning and cognitive development; recent evidence shows that they may be more related than previously thought (Quak et al. 2015).

In a study with an older population (O’Brien et al. 2017), we tested aged 60+ before and after one bout of open- or closed-skill exercise and compared their performance with a control group of participants who took part in a reading club for one session of the same time length. Results showed that immediate memory, assessed with the Digit Span forward test, improved after both exercise types, while benefits on multisensory perception, assessed with the Sound-Induced Flash Illusion (SIFI) (Shams et al. 2000), were found only for the open-skill group.

Both WM and multisensory perception characterise everyday functioning and are impaired in developmental conditions (e.g. ADHD (Klingberg et al., 2002), and dyslexia (e.g. Smith-Spark and Fisk, 2007)). Studies on adult populations report moderate effects of acute exercise on WM performance (Roig et al., 2013). Memory benefits follow an open-skill exercise session (Pesce et al. 2009), yet specific exercises appear to differentially affect memory subcomponents in intervention studies (Alesi et al. 2016).

While memory and executive function represent traditional fields of investigation in relation to exercise, the potential benefits in terms of perceptual abilities are less studied. Nonetheless, efficient multisensory integration allows us to accurately perceive our surroundings and is fundamental to our everyday interactions with the world. Research indicates that multisensory integration remains malleable throughout childhood (Kaganovich 2016; Nava and Pavani 2013), and early adolescence (Brandwein et al. 2010; Chen et al. 2016). Considering the importance of efficient multisensory processing for cognitive development, and the malleability of this process in childhood, it is of relevance to determine whether exercise can represent a way to enhance multisensory integration abilities in children, as it has been found for older adults (Mahoney et al. 2015; Merriman et al. 2015; O’Brien et al. 2017).

In the present study, we built on O’Brien et al. (2017), and we hypothesised that an acute bout of exercise will facilitate cognitive performance in school aged children. Additionally, we expected exercise type to moderate this effect, such that open-skill exercise will lead to greater cognitive benefits compared to other exercise type and the no-exercise group.

## Material and Methods

### Participants

Fifty-one children (28 male) aged between 6 and 8 (*M*= 6.94, *SD*= .54) participated. All participants were typically developing children who reported normal or corrected-to-normal vision and hearing. Participants within each school were randomly assigned to one of three groups; two experimental groups (open-skill exercise, closed-skill exercise) and one control group (no exercise). Each group performed a distinct activity between two experimental testing sessions (pre- and post-test). An additional group of 16 children were tested who undertook the GoNoodle® in class exercise; this group in not included here as children were not randomly assigned to this group.

Those in the open-skill group engaged in open-skill exercises (e.g. basketball, soccer and/or tennis matches), those in the closed-skill group performed exercises classified as closed-skill (e.g. running races, circuit training, skipping) in presence of the other students. The control group engaged in a sedentary classroom activity (i.e. revised homework).

All activities were performed in group and had an approximate duration of 30 minutes. All groups consisted of both male and female participants. See Table 1 for descriptive characteristics for each group.

**Table 1.** Descriptive characteristics for each group

|  | Open-skill | Closed-skill | Control | p value |
| --- | --- | --- | --- | --- |
| *N* | 16 | 16 | 19 |  |
| Age | 7 (0.52) | 6.69 (0.06) | 7 (0.47) | 0.24 |
| Gender | M: 37.5%, F: 62.5% | M: 43.8%, F: 56.3% | M: 78.9%, F: 21.1% | <.001* |
| HR pre | 80.2 (7.85) | 78.83 (7.12) | 77 (6.99) | 0.63 |
| HR post | 123 (12.06) | 125.42 (13.39) | 73.89 (6.68) | <.001* |
| Bout duration | 31.56 (2.39) | 28.75 (4.28) | 27.63 (2.57) | <.001* |
| Handedness | R: 87.5%, L: 6.3%, R/L: 6.3% | R: 93.8%, L: 6.3% | R: 78.9%, L: 21.1% | 0.34 |
| Eyesight | VG 93.8%, Ok 6.3% | VG 87.5%, G 12.5% | VG 100% | 0.28 |
| No. with glasses (%) | 1 (6.2%) | 0 (0%) | 3 (30%) | 0.15 |
| Hearing | VG 100% | VG 75%, G 25% | VG 89.5%, G 10.5% | 0.11 |
| Memory | VG 81.3%, G 18.8% | VG 81.3%, G 12.5%, Ok 6.3% | VG 94.7%, G 5.3% | 0.7 |
| Y-PAQ score<br>(MET minutes) | 2166.57<br>(1524.25) | 1947.33 (536.79) | 2323.16<br>(847.18) | 0.99 |

Ethical approval was granted by the Ethics Committee, University College Cork. All guidelines set out by the Declaration of Helsinki were adhered to. Both children and their parents/guardians provided written informed consent.

### Materials

#### Baseline Measures

Data on participant age, gender, handedness, parental occupation and self-reported health were collected using a self-report questionnaire. To check for exercise intensity across exercise groups, heart rate (HR) was assessed at pre- and post-test. HR was measured using a clinically validated upper-arm blood pressure monitor; OMRON M2 (hem-7117-e; OMRON Healthcare Europe B.V., Hoofdorp, Netherlands), with measurements taken on the non-dominant arm while seated. Where participants reported discomfort with the apparatus, a HR estimate was obtained using a free iTunes app ‘Cardiio’ (Version 3.3.1; Cardio, Inc., 2016), administered using an iPad mini (model A1432). HR scores are expressed as beats-per-minute. The Youth Physical Activity Questionnaire (Y-PAQ) assessed recent physical activity levels (Corder et al. 2009).

#### Verbal Working Memory

Verbal WM was assessed using the Backward Digit Span, a subtest of the WISC-IV (Grizzle 2011). The task involves the verbal presentation of digit series and requires participants to repeat the series in reverse order. Score was calculated as the highest number of correct digits remembered.

#### Visuospatial short-term memory

Participants’ visuospatial WM (i.e. ability to recall object and spatial location) was assessed using a computerised version of the Corsi Block Tapping test (Mueller and Piper 2014), a validated measure (Kessels et al. 2000). The task was downloaded from http://pebl.sourceforge.net/ and administered using a Toshiba Satellite laptop (screen dimensions: 34cm×19cm, resolution: 1366×768, refresh rate: 60 Hz) with maximum screen brightness. Task progression, discontinuation and scoring procedures were identical to those used for the Backward Digit Span.

#### Motor Short-term Memory

Participants’ ability to recall information on motor actions was assessed using a motor span task, whereby all the stimuli consisted of meaningless gestures selected among the proximal meaningless gesture of the STIMA test for apraxia (Tessari et al. 2015). The task requires participants to remember and reproduce a sequence of upper-arm movements performed by the experimenter (each movement lasted one second) in the order in which they were presented. After the execution of each movement the experimenter returned to a neutral position with the arms at her side. Task progression, discontinuation and scoring procedures were identical to the previous memory tasks.

#### Multisensory Perception

The Sound-induced Flash Illusion, SIFI (Shams et al. 2000, 2002) was used to assess participants’ multisensory integration efficiency. The SIFI consists of the illusory perception of two flashes (dots) when one flash is presented simultaneously with two beep sounds.

Susceptibility to the SIFI is considered an indicator of efficient multisensory processing and has been used in this capacity in studies with preadolescents (Innes-Brown et al. 2011; Nava and Pavani 2013; Tremblay et al. 2007). SIFI is also associated with efficient cognitive processing in older adults (Chan et al. 2015; Hernández et al. 2019), although its association with WM in children has not been tested to date.

In the SIFI the flash is constituted by a dot (1.5 cm diameter) presented 5 cm below fixation (duration 16ms) and the auditory beep has duration 10ms (1ms ramp, 3500Hz), transmitted via the computer loudspeakers. The task includes unisensory conditions (1 flash, 2 flashes; 1 beep, 2 beeps) and multisensory conditions; the multisensory conditions are either congruent (1flash/1beep, 2flashes/2beeps) or incongruent (1flash/2beeps). In the 1flash/2beeps and 2flashes/2beeps conditions, the stimulus Onset Asynchrony (SOA) varied amongst trials, with SOAs of 70, 110, 150 and 230ms. The software suite Presentation (version 18; www.neurobs.com) was used to programme and administer the SIFI. The SIFI was administered on a laptop (see above). Volume and brightness of the laptop were set to maximum to ensure consistent presentation of stimuli for all participants.

Participants were administered two blocks of trials, in the first block, comprising visual and audio-visual stimuli, participants were asked to report how many flashes they saw. They were instructed to ignore the beeps and only count flashes. The second SIFI block consisted of only auditory stimuli to ensure participants could hear the beeps. For these trials, participants were asked to report the number of beeps heard. Participants’ responses were verbal and the experimenter inputted them using the laptop keyboard. The SIFI task comprised of 92 trials; 38 unisensory trials and 54 multisensory trials. The 54 multisensory trials comprised 6 one flash with one beep, 24 two flash with two beeps (6 per SOA) and 24 illusory trials (i.e. one flash with two beeps; 6 per SOA). Scores were calculated as the mean proportion of correct responses to each trial type. The duration of the test is approximately 6-8 minutes.

### Procedure

Participation involved completing two testing sessions, one before (pre-test) and one immediately after an activity bout (post-test). Participants were tested individually. At pre-test, resting HR was measured and questionnaires were administered followed by the four cognitive tasks (i.e. three memory tasks and the SIFI). Participants then completed an activity session corresponding to their assigned group. Post-test, immediately after activity cessation, consisted of repeating the HR measurement and cognitive tasks. Cognitive task administration was counterbalanced to minimise order effects. Pre- and post-test lasted approximately 20 minutes and 12 minutes respectively.

### Analysis

Statistical analysis was conducted using IBM SPSS (version 22). Conformity to parametric assumptions can be assumed unless violations are specified. The alpha level was *p*< .05. Effect sizes are reported in partial eta squared 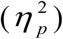 and interpreted accordingly; <0.08 (small), 0.08 - 0.14 (medium) and >0.14 (large).

#### Working Memory Analyses

Group (open-skill, closed-skill, control) and Time (pre, post) were entered as independent variables, with dependent variables being memory span. A mixed measures analyses of variance (ANOVA) was conducted on memory spans (i.e. verbal, visuospatial and motor working memory). Significant interactions were analysed using Bonferroni corrected tests.

#### Multisensory Perception

Dependent variables were *d’* scores, which reflects perceptual sensitivity and performance on the SIFI task. *d’* capture a participant’s ability to correctly perceive 2 flashes when they are real (i.e. correctly detecting 2 flashes in 2flash/2beeps conditions) relative to their susceptibility to the illusion (i.e. incorrectly perceiving 2 flashes in 1flash/2beeps conditions). Sensitivity was calculated using the following formula: d’ [z(hits) – z(false alarms)], where hits were proportion of correct responses on 2flash/2beep conditions and false alarms were proportion of incorrect responses to 1flash/2beep conditions. and z refers to the inverse cumulative norm (Green and Swets 1966).

## Results

No baseline group differences were present for age, Y-PAQ scores, HR or self-report variables, while both exercise groups displayed significantly higher post-test HR relative to the control group, while it did not differ between exercise groups (see Table 1). Duration of activity bout (in minutes) significantly differed across groups [*H*(2)= 11.38, *p*<.01], with the control group registering shorter activity bouts compared to the open-skill group (see Table 1). Given the documented association between exercise duration and cognitive outcomes post-exercise (McMorris et al. 2011), even if the difference was only few minutes, bout duration was entered as a covariate in subsequent analysis procedures to control for these group differences.

### Working Memory

A 2 (Time) × 3 (Memory Task) × 3 (Group) Mixed ANCOVA was conducted on span scores. Bout duration was entered as covariate, Time (pre, post) and Memory Task were the within-subjects factors and Group (open-skill, closed-skill, control) the between-subjects factor. Table 2 displays mean scores per group for each of the memory tasks.

**Table 2.** Mean span scores on memory tasks (standard error in parentheses)

|  | Motor | Verbal | Visuospatial |
| --- | --- | --- | --- |

|  | T1 | T2 | T1 | T2 | T1 | T2 |
| --- | --- | --- | --- | --- | --- | --- |
| Open-skill | 3.31 (0.19) | 3.5 (0.18) | 3.06 (0.15) | 3.63 (0.15) | 4.44 (0.19) | 4.67 (0.19) |
| Closed-skill | 3.25 (0.19) | 4 (0.18) | 2.94 (0.25) | 3.13 (0.15) | 4.06 (0.19) | 4.44 (0.19) |
| Control | 3.58 (0.18) | 3.68 (0.17) | 3.26 (0.14) | 3.37 (0.14) | 4.32 (0.17) | 4.32 (0.17) |

For memory span scores, the ANCOVA revealed a significant Time × Group interaction effect [*F*(2, 94) = 3.99, *p* = .025, 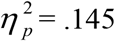]. A significant three-way interaction Time × Group × Memory Task emerged [*F*(3.76, 88.48) = 2.63, *p* = .043, 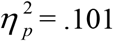]. No other main or interaction effects were significant. Post hoc analysis of the three-way interaction was performed by three separate paired samples t-test for each group, comparing scores at pre- to post-test. This analysis revealed that, when Bonferroni correction was applied, the open-skill group showed a positive effect of exercise on verbal working memory (*t*(15) = −3.09, *p* = 0.007). The closed-skill group improved in motor span (*t*(15) = 3.97, *p* = 0.002). The control group did not show any significant effect (see Figure 1a and Figure 1b).

**Fig 1a.**
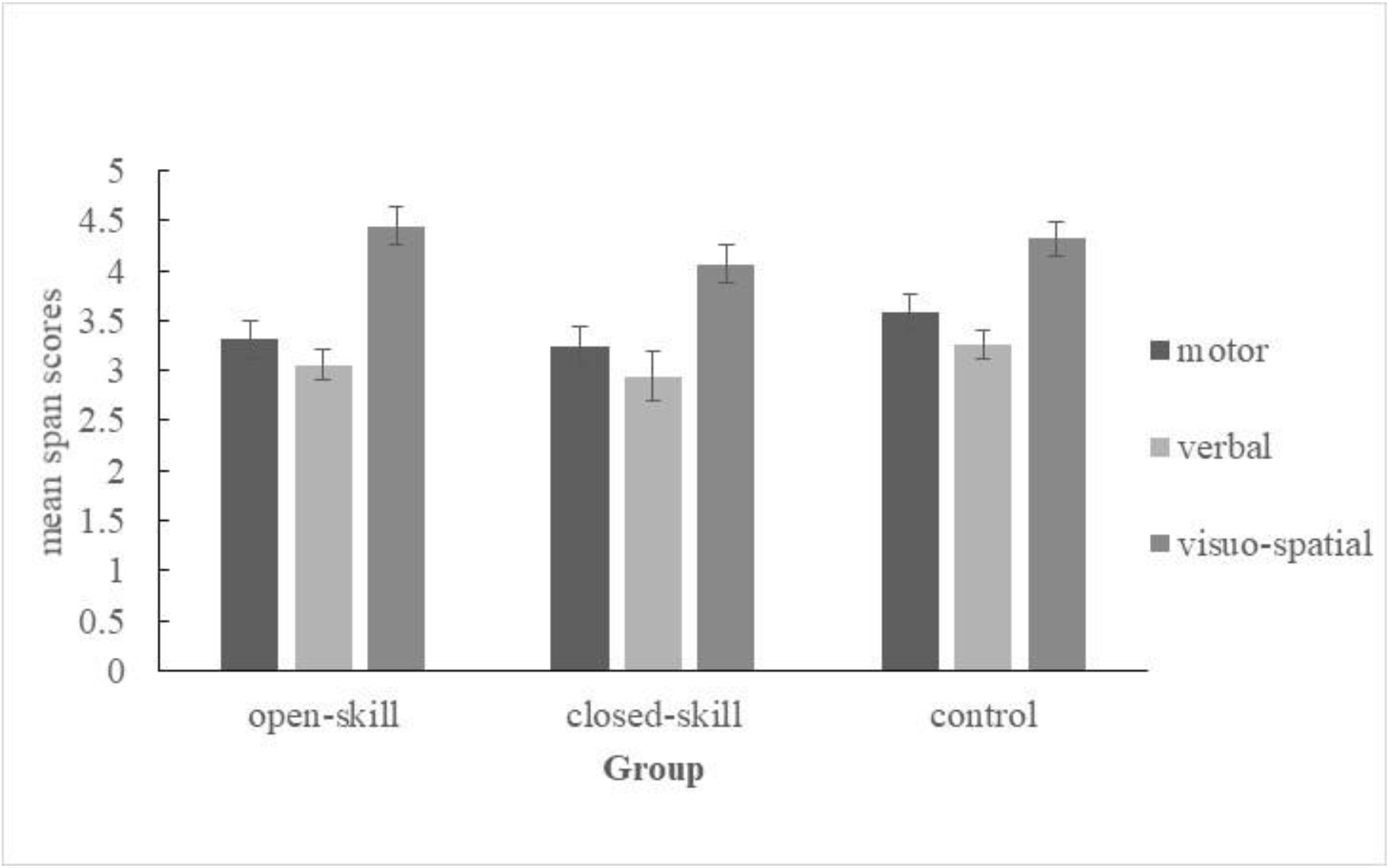
Mean span scores at Time 1 (error bars indicate one standard error of the mean)

**Fig 1b.**
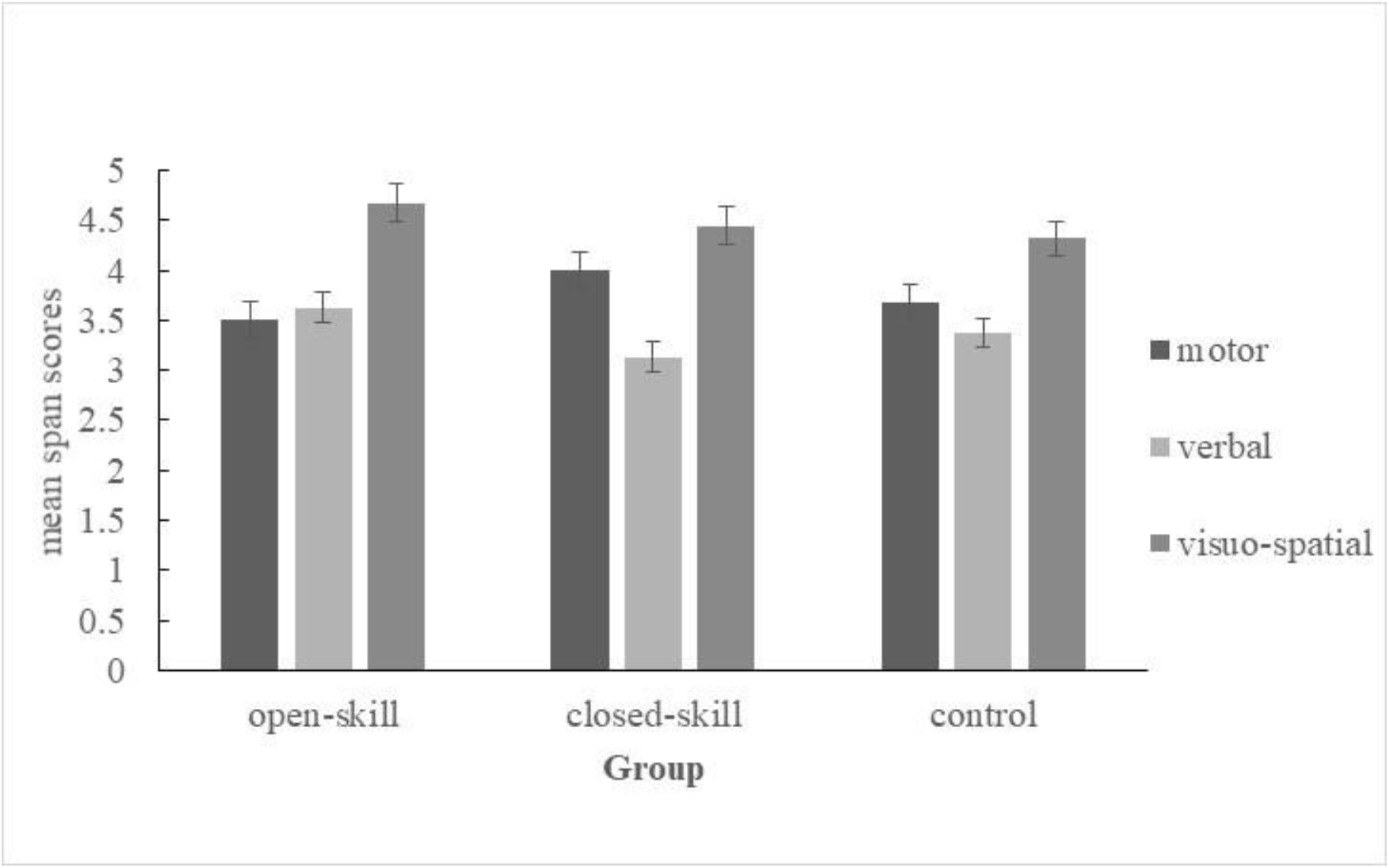
Mean span scores at Time 2 (error bars indicate one standard error of the mean)

### Multisensory Perception

The sample for this analysis was 46 due to removal of participants who always pressed the same key or did not complete the task. Characteristics of this sub-sample are presented in Appendix A. Independent variables for this analysis were Group, Time and SOA (70, 110, 150, 230ms). Preliminary analyses revealed participants perceived the illusion and no baseline group differences in susceptibility to the illusion existed: a Mixed ANOVA with Group as between participants factor and SOA (70, 110, 150, 230ms) and Condition (illusory (1beep/2flashes, congruent 2beeps/2flashes) as within participants factors, was run on participants’ baseline SIFI data (t1). Relevant to our aims, a main effect of Condition (*F*(1,43) = 382.99, *p*<.001, 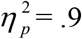) was found, with participants displaying significantly more correct responses to congruent conditions (*M*= .92, *SD*= .17) compared to illusory conditions (*M*= .16, *SD*= .2), confirming participants experienced the illusion. Of note, a main effect of Group did not emerge [*F*(2, 43)= .65, *p*= .53, 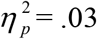], with groups performing comparably on baseline conditions.

#### Perceptual Sensitivity Analysis

An analysis procedure based on Signal Detection Theory was conducted to explore whether exercise had an effect on participants’ perceptual sensitivity (i.e. ability to detect 2 real compared to 2 illusory flashes. Dependent variables for this analysis (*d’* scores) met all assumptions for parametric tests (following Box-Cox transformations).

A Mixed ANCOVA was performed on *d’* scores, with Time, Group and SOA entered as factors. Bout duration served as covariate. There was no effect of Time or SOA on *d’* scores. A significant main effect of Group was observed [*F*(2, 42) = 6.75, *p* = .003, 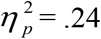]. A significant Time × Group interaction effect also emerged [*F*(2,42) = 5.55, *p* = .007, 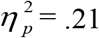]. To follow-up on the significant interaction, a series of paired sample t-tests were performed, with Time as the factor. The open-skill group significantly improved in *d’* scores from pre (*M*= .46, *SD*= .46) to post-test (*M*= 1.14, *SD=*. 68), *t*(14)= −3.17, *p*= .007, *d*= 1.17. The closed-skill group also significantly improved in sensitivity, *t*(13)= −3.02, *p*= .01, *d*= .87, from pre (*M*= .61, *SD*= .45) to post-test (*M*= .99, *SD*= .42). By contrast, sensitivity scores for the control group (*t*(16)= 1.29, *p*= .22, *d*= .42) did not significantly differ between testing sessions (pre, *M*= .29, *SD*= .92; post-test, *M*= -.068, *SD*= .79). Figure 2 depicts this significant Time × Group interaction effect. In order to ensure that these differences were not solely due to HR, we introduced both duration of bout and HR post exercise as covariates, which led to a significant interaction (Time × Group × SOA, *F*(6,105) = 2.77, *p* = .016, 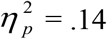), indicating that the difference between exercise groups and control groups was not solely due to the difference in heart rate achieved after the different activities (proportion of correct responses is reported in Appendix B).

**Fig 2.**
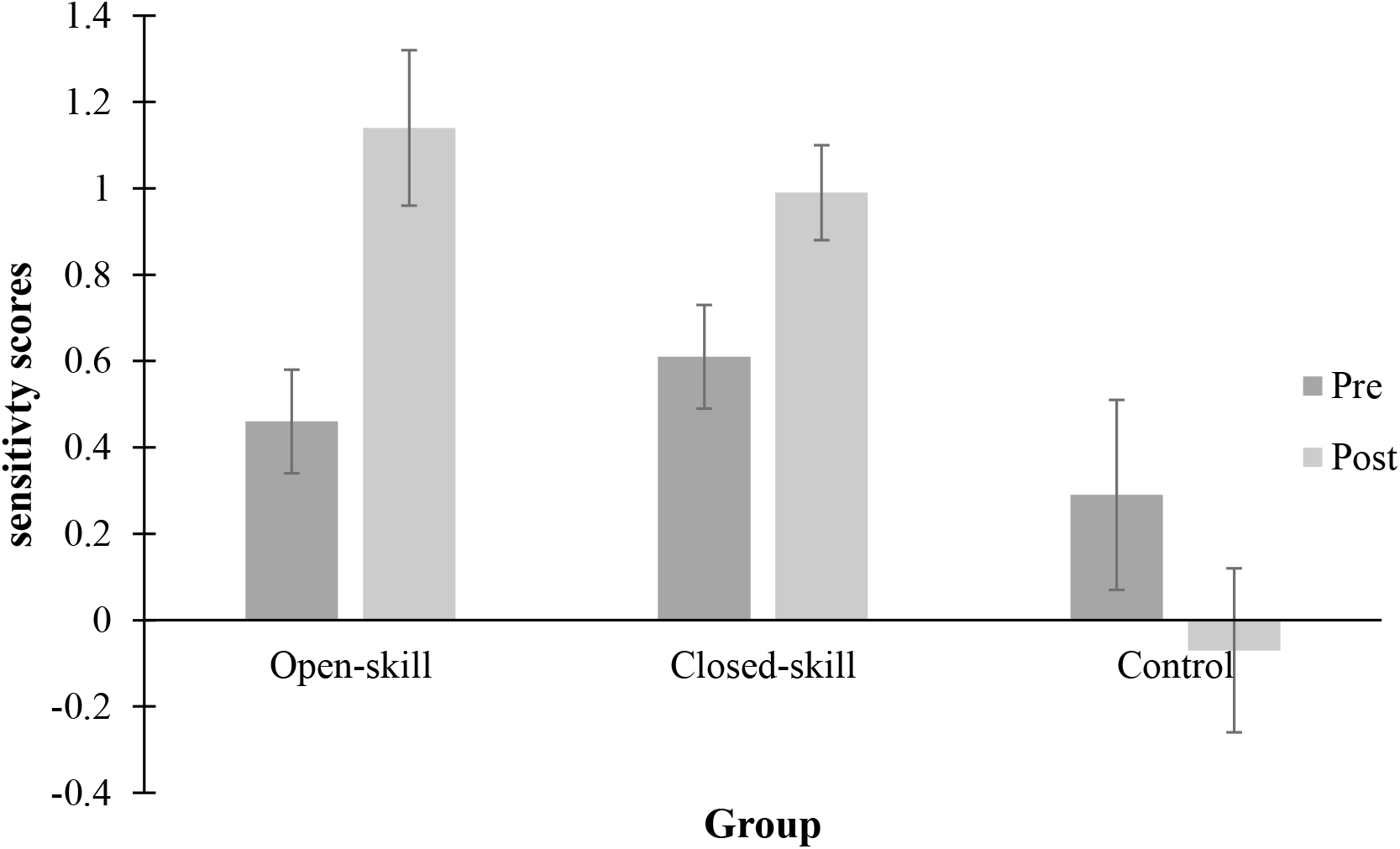
Time x Group interaction on *d’* scores (error bars indicate one standard error of the mean)

As recent evidence indicates that multisensory perception could be associated with WM capacity, we conducted an exploratory paired correlations between the WM scores and the *d*’ (SIFI sensitivity). No significant correlations were found between any of the WM and d’ pre, nor post exercise or control activity.

## Discussion

There is a lack of understanding of the benefits, or lack of thereof, of one bout of open- or closed-skill exercise in school-aged children. We focused on two cognitive functions, multisensory perception, i.e. the SIFI, and WM, which are crucial in supporting cognition and function. Specifically, the primary aim of this research was to examine the potential mediating role of exercise type on the effect of acute exercise in line with what we found for older adults (O’Brien et al. 2017).

Firstly, exercise type did emerge as a reliable moderator of the exercise-cognition relationship in the present study. In accordance with the cognitive engagement hypothesis, it was expected that exercise type would modulate the effect of exercise on WM, such that open-skill exercise would produce the most cognitive benefits. A session of open-skill exercise had a positive effect on digit span; on the other hand, closed-skill exercise improved motor short-term memory (i.e. only a sub-component/slave-system of working memory according to Baddeley’s model; e.g. (Baddeley 2011)). No benefits were found for visuospatial short-term memory. For open skill exercise, results are in line with literature often reporting a beneficial effect of intermediate-intensity exercise on many working memory tasks (see McMorris et al. (2011) for a recent meta-analysis) and they are novel, as the task used here (backword digit span) is less heavily dependent on central executive compared to others previously used (e.g. switching task, Stroop, flanker task). As to motor WM, which was improved with closed-skill exercise, it is clear that the motor aspect of memory would be most apt to improve after exercise, as opposed to classroom work, given the invariable need to process and reproduce motor movements during physical activity, however why this is specific to closed-skill remains an open question.

It is worth noting that the duration of the exercise session here is approximately 30 minutes, which is an intermediate dose of exercise, as opposed to a high dose, see (Tomporowski et al. 2015), therefore it suggests that benefits can be found with half an hour of targeted physical activity, which is promising in terms of applicability to a variety of school settings.

Findings in relation to multisensory perception are consistent with expectation and existing research (Innes-Brown et al. 2011). Firstly, results indicated 6-8 year old children are susceptible to the SIFI and furthermore, displayed a high degree of susceptibility, with participants on average experiencing illusions 84% of the time. This suggests children are unable to ignore task irrelevant stimuli (i.e. the beeps) and hence integrate extensively, a pattern similar to that observed for older adults with the SIFI task (Setti et al. 2011). This finding is in line with studies reporting that children exhibit less fine-tuned multisensory integration in childhood (Adams 2016) and adolescence (Brandwein et al. 2010). Supporting our hypothesis that open-skill exercise would facilitate an environment conducive for perceptual training, participants in the open-skill group displayed improved perceptual sensitivity post-exercise. However, the closed-skill group also showed improved perceptual sensitivity. These results contrast with our study in older adults (O’Brien et al. 2017), which reported only open-skill exercise led to immediate benefits for multisensory perception. The difference could be methodological, as in O’Brien et al. (2017) the open- and closed-skill groups also differed in exercise intensity, unlike the current study (as shown by lack of HR differences). It is also notable that the predictions of the present study, being the first to explore acute exercise and multisensory perception in children, were based on research with adults. With both open- and closed-skill exercise conferring a perceptual benefit in children, this suggests that it could be the physiological arousal more than cognitive engagement aspect of exercise that led to perceptual benefits in the present study; however HR do not significantly impact on the results when added as covariate, therefore more research is needed.

The finding that either a session of open-skill or closed-skill exercise can improve multisensory perception in schoolchildren opens up numerous research avenues with potential applied implications. Considering multisensory perception is impaired in several developmental disorders, such as dyslexia (Hahn et al. 2014), and autism (Baum et al. 2015) and with few viable interventions avenues currently available, exercise represents a potential intervention to train multisensory perception. Furthermore, as both open- and closed-skill exercises are already widely used in schools (Masini et al. 2019) and popular amongst children, the use of exercise as an interventional strategy for perceptual training is a potentially attractive and cost-effective option.

Limitations of the present study must be considered. Firstly, while we endeavoured to randomise the sample, in the present study only partial randomisation was achieved, based on school and class availability. In addition, it is recommended that future research studies using the SIFI paradigm adopt larger SOAs as the SOAs used in the current study (i.e. 70, 110, 150, 230ms) were possibly too brief to capture a comprehensive estimate of children’s multisensory temporal binding window, at this developmental stage.

Secondly, the association between habitual exercise and benefits derived from the one bout has been limited in the present study by the use participant self-report for past week physical activity (YPAQ), which is bound by the apparent low reliability in this young sample. The case has been made that children under the age of 10 do not possess the required cognitive resources to accurately recall their recent physical activity patterns (Baranowski 1988). For future research, an objective measure of recent physical activity (e.g. an accelerometer) should be favoured. An unexpected technical issue in relation to the heart rate assessments also limits our results. As some children were uncomfortable with the use of a digital blood pressure monitor to assess heart rate, an iTunes app was used to gain a proxy measurement. With no reliability data available for the app in children, the accuracy of these heart rate readings cannot be determined. Validation studies of these apps are needed, particularly for children, where the use of clinical equipment may pose difficulties. Despite these limitations, which primarily relate to the measurement of control rather than experimental variables, the current study provides novel insights into the exercise-cognition relationship.

Noteworthy strengths of this study include the simultaneous exploration of a range of cognitive functions, the use of a suitable control group (i.e. cognitively engaging sedentary activity; homework corrections), and its ecological validity (i.e. conducted in school setting). This study contributes to the mounting evidence for the selective nature of the exercise-cognition link and presents novel findings on the benefits of a classroom activity break.

## Declarations

The authors have no specific funding source to declare, the research was part of the Masters in Applied Psychology by JOB supervised by AS. The authors declare no conflicts of interest in relation to this research. None of the authors declare competing financial interests. No formal consent was obtained for lodging the data on a public repository, therefore the data, in semi-anonymised format, can be obtained from JOB or AS. Authors contribution: JOB, AS, GO, AT designed the study; JOB collected and analysed the data; JOB and AS drafted the manuscript; GO, AT revised the manuscript.

## Appendix A

**Table 1A.** Descriptive Characteristics by Group for the SIFI test

|  | Open skill | Closed skill | Control | *p*<br>value |
| --- | --- | --- | --- | --- |
| *N* | 15 | 14 | 17 |  |
| Age (in years)<sup>a</sup> | 7 (0.54) | 6.64 (0.63) | 6.94 (.43) | 0.17 |
| Gender | Male 40%, Female 60% | Male 42.9%, Female 57.1% | Male 76.5%, Female 23.5% | 0.07 |
| HR pre<sup>a</sup> | 80.79 (7.8) | 80.4 (6.33) | 77.31 (7.36) | 0.38 |
| HR post<sup>a</sup> | 124.43 (11.12) | 126.2 (13.42) | 74.06 (6.97) | <.001* |
| Bout duration<sup>a</sup> | 31.33 (2.29) | 29.29 (4.32) | 27.65 (2.58) | .007* |
| Handedness | R: 86.7%,<br>L: 6.7%,<br>R/L: 6.7% | R: 92.9%,<br>L: 7.1% | R: 88.2%,<br>L: 11.8% | 0.94 |
| Self-report Eyesight | VG 93.3%,<br>Ok 6.7% | VG 92.9%,<br>G 7.1% | VG 100% | 0.39 |
| Self-report Hearing | VG 100% | VG 85.7%, G 14.3% | VG 94.1%, G 5.9% | 0.1 |
| Self-report Memory | VG 80%, G 20% | VG 85.7%, G 14.3% | VG 94.1%, G 5.9% | 0.48 |
| Y-PAQ (MET-mins)<sup>a</sup> | 2230.04 (1561.08) | 2007.08 (554.21) | 2262.59 (872.97) | 0.79 |

*Note.* <sup>a</sup> denotes scores expressed as the mean value with standard deviation in parentheses. HR, heart rate; Y-PAQ, youth physical activity questionnaire; VG, very good; G, good. P value refers to results of One Way ANOVA with group entered as a factor, or chi-square analyses in the case of categorical variables. \* denotes significant *p* values.

## Appendix B

**Table 1B.** Proportion of correct responses in illusory trials in the SIFI test

|  | Pre test |  |  |  | Post test |  |  |  |
| --- | --- | --- | --- | --- | --- | --- | --- | --- |
|  | 1f2b 70 | 1f2b 110 | 1f2b 150 | 1f2b 230 | 1f2b 70 | 1f2b 110 | 1f2b 150 | 1f2b 230 |
| Open skill | 0.22 (0.04) | 0.08 (0.03) | 0.13 (0.04) | 0.11 (0.04) | 0.25 (0.04) | 0.13 (0.03) | 0.21 (0.07) | 0.33 (0.05) |
| Closed skill | 0.11 (0.04) | 0.11 (0.03) | 0.04 (0.02) | 0.23 (0.05) | 0.27 (0.04) | 0.11 (0.04) | 0.11 (0.04) | 0.18 (0.04) |
| Control | 0.34 (0.08) | 0.22 (0.08) | 0.24 (0.08) | 0.26 (0.09) | 0.15 (0.04) | 0.13 (0.07) | 0.19 (0.08) | 0.24 (0.08) |

